# FiGS-MoD: Feature-informed Gibbs Sampling Motif Discovery Algorithm for Mapping Human Signaling Networks

**DOI:** 10.1101/2025.09.26.678911

**Authors:** Yitao Sun, Yu Xia, Jasmin Coulombe-Huntington

## Abstract

**Motivation:** Short linear motifs (SLiMs) are short sequence patterns that mediate transient protein-protein interactions, often within disordered regions of proteins. SLiMs play central roles in signaling, trafficking, and post-translational regulation, but their short length and low complexity make them difficult to identify both experimentally and computationally. Since the latest release of motif discovery tools like MEME Suite, the availability of protein-protein interaction data (e.g., BioGRID) has increased by more than fivefold, providing richer network contexts where SLiMs can be inferred from recurring patterns of interaction. Combined with recent advances in machine learning, this creates new opportunities for large-scale, high-resolution motif discovery.

**Results:** We present FiGS-MoD, a **F**eature-**i**nformed **G**ibbs **S**ampling **Mo**tif **D**iscovery algorithm with two key innovations: (i) incorporating biased sampling informed by residue-level features, including Protein Language Model (PLM) embeddings, AlphaFold2-derived disorder and solvent accessibility, and evolutionary conservation, and (ii) replacing the traditional position-specific scoring matrix (PSSM) with a Hidden Markov model (HMM) to accommodate insertions and deletions. We applied our algorithm to 12,765 sub-networks from the human interactome and provided 221,840 human SLiM predictions and quality scores as a public resource, along with the tool itself. Our method outperformed MEME in terms of recovering known motifs from the Eukaryotic Linear Motif (ELM) database and phosphosites from PhosphoGRID. Through three case studies, we further highlight the biological relevance of our results and the generalizability of the method to diverse motif classes.

**Availability and Implementation:** Source code and a predicted SLiM dataset using FiGS-MoD are freely available at https://github.com/Eric3939/FiGS-MoD

## 1 Introduction

Short linear motifs are short stretches of amino acids, typically 3 to 11 residues, embedded within protein sequences, often located in intrinsically disordered regions (IDRs) (Davey et al., 2012). Despite their brevity and low information content, SLiMs are critical for a variety of cellular processes, including protein localization, modification, and transient signaling interactions. Examples include SH3-binding motifs, phosphorylation sites, and PDZ-binding domains (Ren et al., 2008; Sologova et al., 2022; Van Roey et al., 2014).

A motif class represents a general pattern (often described via regular expressions, PSSMs, or HMMs), while a motif instance refers to a specific occurrence of the motif in a protein, typically localized to a defined sequence position. The ELM database currently catalogs over 350 motif classes and ∼4300 validated motif instances (Kumar et al., 2024).

*De novo* SLiM discovery is a challenging research question. Experimental discovery of SLiMs remains difficult. Techniques like yeast two-hybrid or affinity capture are less effective for the weak and transient interactions that SLiMs typically mediate (Davey et al., 2023). Structural methods (e.g., cryo-EM or X-ray crystallography) often exclude disordered regions, further limiting the ability to resolve motifs.

Consequently, computational methods have become central to SLiM discovery (Edwards & Palopoli, 2014). Traditional approaches like MEME search for overrepresented motifs within protein sets—often proteins that interact with a common partner—using techniques such as expectation-maximization or Gibbs sampling (Bailey et al., 2015).

There are two major reasons why it is timely to revisit computational SLiM discovery: The first is the massive expansion of public datasets. Since the original publication of MEME, protein-protein interaction (PPI) databases like BioGRID have grown significantly. Namely, BioGRID documented ∼190,000 human interactions in 2015, compared to over 1 million interactions today which is a more than five-fold increase (Oughtred et al., 2021). Second, recent advances in machine learning, particularly PLMs and structure prediction tools like AlphaFold, provide residue-level annotations of disorder, accessibility, and functional information. These features can now be used to guide motif discovery in ways that were not previously possible.

There are two limitations of previous computational methods. The first is the motif pattern modeling using PSSMs or their variants. While efficient, PSSMs assume fixed-length motifs and do not model insertions or deletions, which are common in flexible, disordered regions: 86 out of 288 ELM motif classes (29.9%) contain one or more indels. In contrast, HMMs naturally model gapped alignments and have been highly successful in protein family classification (e.g., via HMMER) and multiple sequence alignment (e.g., CLUSTAL) (Finn et al., 2011; Sievers et al., 2011). However, HMMs have not been widely adopted for de novo motif discovery in the Gibbs sampling framework. The second limitation is a lack of biological data (e.g., structural information and evolutionary information) in the discovery tool, as most tools are sequence-based only.

In this work, we present FiGS-MoD, a novel feature-informed Gibbs sampling algorithm for *de novo* SLiM discovery. By incorporating residue-level priors from protein language models, AlphaFold2-derived disorder and solvent accessibility, and evolutionary conservation, and replacing the traditional PSSM scoring with profile HMM, FiGS-MoD guides sampling toward residues more likely to harbor functional motifs. We scanned 12,765 PPI sub-networks from BioGRID for potential SLiMs bound by a common interactor protein and provided all predictions and their quality scores as a public resource. We benchmark FiGS-MoD against MEME, a widely used motif discovery algorithm, using ELM and BioGRID, showing improved recovery of known motifs and phospohsites, and provide case studies illustrating its biological utility.

## 2 Method

In overview, we first grouped proteins by common interactors, forming 12,765 sub-networks from BioGRID, henceforth referred to as “star” networks (e.g., all partners of a common protein, which serves as the hub node). Residues were annotated with prior probabilities derived from protein language model embeddings, AlphaFold2 structural features (disorder and solvent accessibility), and multiple sequence alignment-based conservation. These priors bias Gibbs sampling by influencing the likelihood of residue selection. Sampling windows of variable length were drawn to construct HMM-based motif models, and acceptance or rejection was guided by motif-model similarity. A dropout strategy further removed poorly fitting windows to mitigate noise in biological data and the dilution of SLiM-mediated interactions with other modes of interactions, including indirect interactions. Predictions were collected after 3,000 iterations, which consistently yielded convergence. An overview of the workflow is shown in Figure 1.

**Figure 1.**
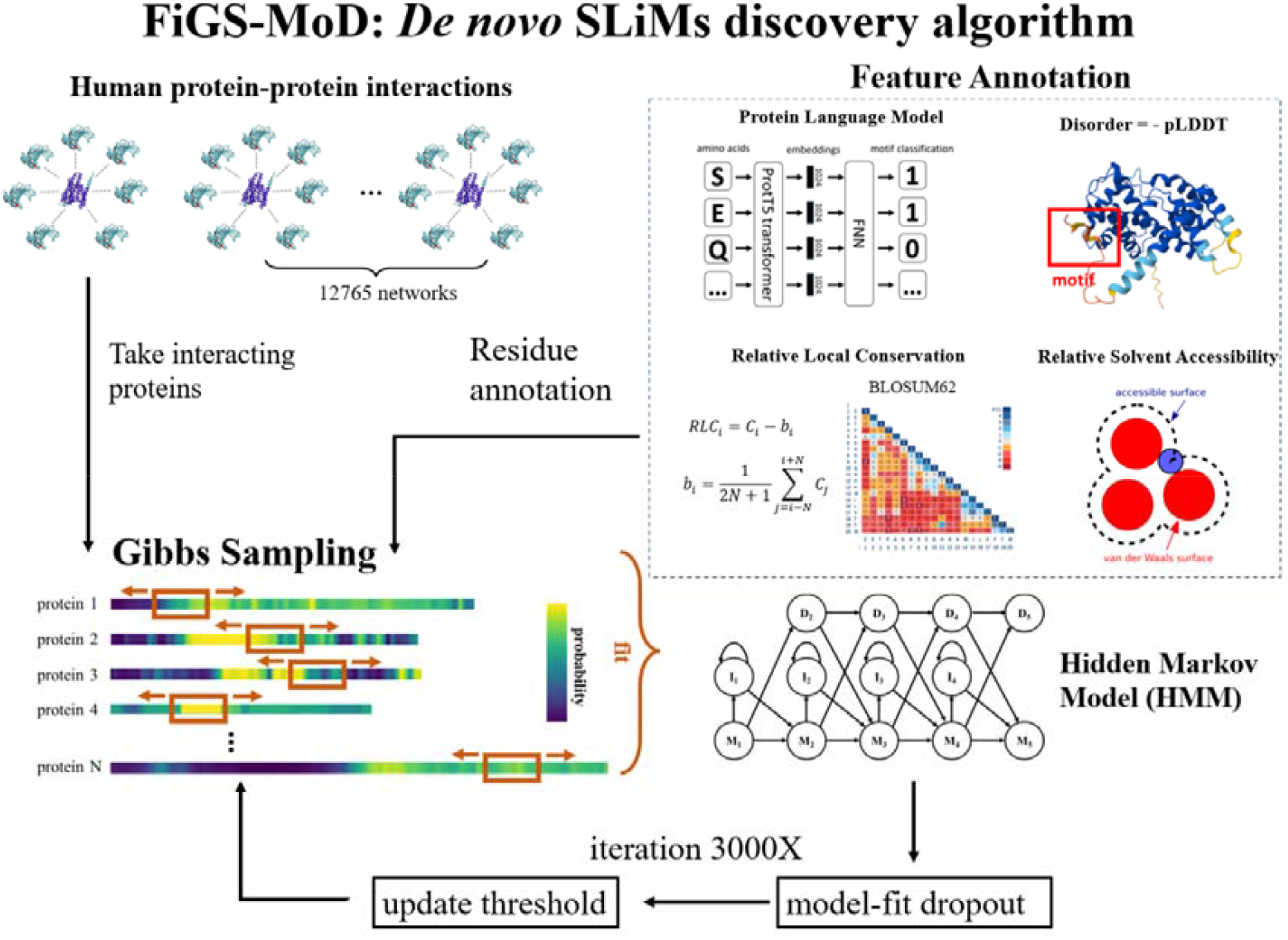
Overview of the FiGS-MoD algorithm for linear motif discovery. Protein-protein interaction data are first grouped into star networks, each consisting of one protein and all of its direct interactors. These star networks serve as the input for Gibbs sampling. Each protein sequence is annotated at the residue level using features from protein language models, AlphaFold2-derived disorder and solvent accessibility, and multiple sequence alignment-based relative local conservation (RLC). These annotations provide priors that bias Gibbs sampling towards residues more likely to contain functional motifs. Sampling windows of variable length are drawn and scored using a profile hidden Markov model (HMM), allowing insertions and deletions. A dropout strategy removes poorly fitting windows to reduce the noise of HMM from non-motif regions. Based on empirical testing, ∼3000 iterations were sufficient for convergence, after which predicted motifs were compiled into a proteome-wide database available on our GitHub repository. As Gibbs sampling is inherently stochastic and does not guarantee a global optimum, repeated runs with different initializations would be required to yield the best possible results.

### 2.1 Feature Annotation

#### 2.1.1 Protein Language Model ProtT5

We used ProtT5 residue-level embeddings from UniProt and trained a three-layer neural network (512→256→128) to classify residues as motif/non-motif using ELM as labels (Elnaggar et al., 2021). Training used an 80/20 split and balanced sampling (16249 residues per class). Test performance achieved 84.8% sensitivity and 84.6% specificity. The sigmoid output provides the final PLM score for each residue. There are 11405811 residues in total in the human proteome, where 16249 residues are positive labels (residues being part of the ELM motifs).

#### 2.1.2 AlphaFold

Two features from the AlphaFold 2 database (Jumper et al., 2021; Varadi et al., 2024) were extracted: disorder and solvent accessibility. The pLDDT has been shown to be a strong indicator of disorder (Zhao et al., 2023), which is derived from the negation of the per-residue pLDDT (pLDDT <50 indicates disorder). The missing data (602 proteins, 3% of the total) were compensated using IUPred3 (Erdős et al., 2021), which is one of the latest sequence-based disorder prediction tools. Disorder scores from IUPred (0-1 range) were rescaled to the 0-100 scale of pLDDT by computing 100×(1-IUPred), ensuring compatibility. Solvent accessibility was computed via DSSP on AlphaFold2-predicted coordinates (Kabsch & Sander, 1983).

#### 2.1.3 Relative Local Conservation

We calculated residue-wise relative local conservation scores (RLCs) for all human proteins. Protein multiple sequence alignments (MSAs) from the UCSC Genome Browser, based on 99 vertebrate genomes, were used as the raw data source (Perez et al., 2025). Compared with earlier approaches, this dataset provides greater phylogenetic depth for conservation analysis.

Because the UCSC human protein sequences do not always match the UniProt reference, we mapped UCSC sequences to UniProt using BLASTp with an identity threshold of ≥90% (Camacho et al., 2009). When a match was found, the UCSC human sequence was replaced with the corresponding UniProt sequence. The resulting MSAs thus contained UniProt human sequences aligned with vertebrate orthologs from UCSC. In practice, 97.7% of mappings were at 100% identity, with nearly all others ≥95%, ensuring that conservation scores were derived from the correct protein sequences. Final MSAs were refined with ClustalO (Sievers et al., 2011).

Conservation was quantified using the BLOSUM62 substitution matrix by comparing each vertebrate residue to the human reference residue (Henikoff & Henikoff, 1992):

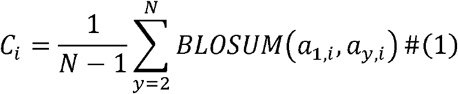

where a_1,j_ is the human residue at position i, a_y,i_ is the corresponding residue from species y, and N is the number of aligned species.

We then derived the relative local conservation (RLC) score, a normalized measure proposed to better capture SLiMs (Davey et al., 2009). Since SLiMs are typically short, conserved stretches embedded within otherwise disordered regions, RLC emphasizes local deviations from the background conservation level:

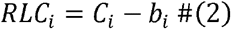

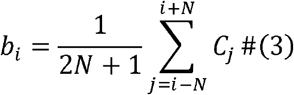

with b_i_ defined as the mean conservation within a 30-residue sliding window centered on residue i.

The four feature scores (ProtT5, disorder, RSA, and conservation) were first z-normalized independently to ensure comparability across scales. These normalized values were then combined into a single score using a weighted linear combination, with ProtT5 assigned a weight of 5 and the other features each assigned a weight of 1, a scheme that performed well in practice. The resulting combined scores were subsequently normalized to sum to 1, defining a categorical probability distribution over residues from which windows were sampled during Gibbs sampling.

### 2.2 HMM-integrated Gibbs sampling

Classical motif discovery algorithms employ Gibbs sampling with PSSMs, but such models cannot account for insertions or deletions. We extended this framework by replacing the PSSM with a profile HMM implemented in Pomegranate (v0.15.0) (Schreiber, 2017), following the design of Durbin’s profile HMM (Durbin et al., 1998). The model included match, insertion, and deletion states for each position. Windows’ similarity to the HMM was quantified as the log-probability under the HMM, and parameters were estimated by Viterbi learning, which provided comparable accuracy to Baum-Welch with reduced runtime.

To accommodate flexible motif lengths, window sizes were sampled from a normal distribution centered on the expected motif length (e.g., N(7, 0.7) for motifs of ∼7 residues). We further introduced a model-fit dropout strategy to relax the assumption that all sequences contain a motif. After each iteration, windows with log probability more than two standard deviations below the mean of the past 100 iterations were masked, ensuring that model updates were driven only by high-scoring candidates. The algorithm is shown in Figure 2 (A), with information content and log probability during training shown in Figure 2 (C) and (D). Additionally, we note that convergence of Gibbs sampling does not guarantee recovery of the global optimum, and thus results may vary across runs with different initializations.

**Figure 2.**
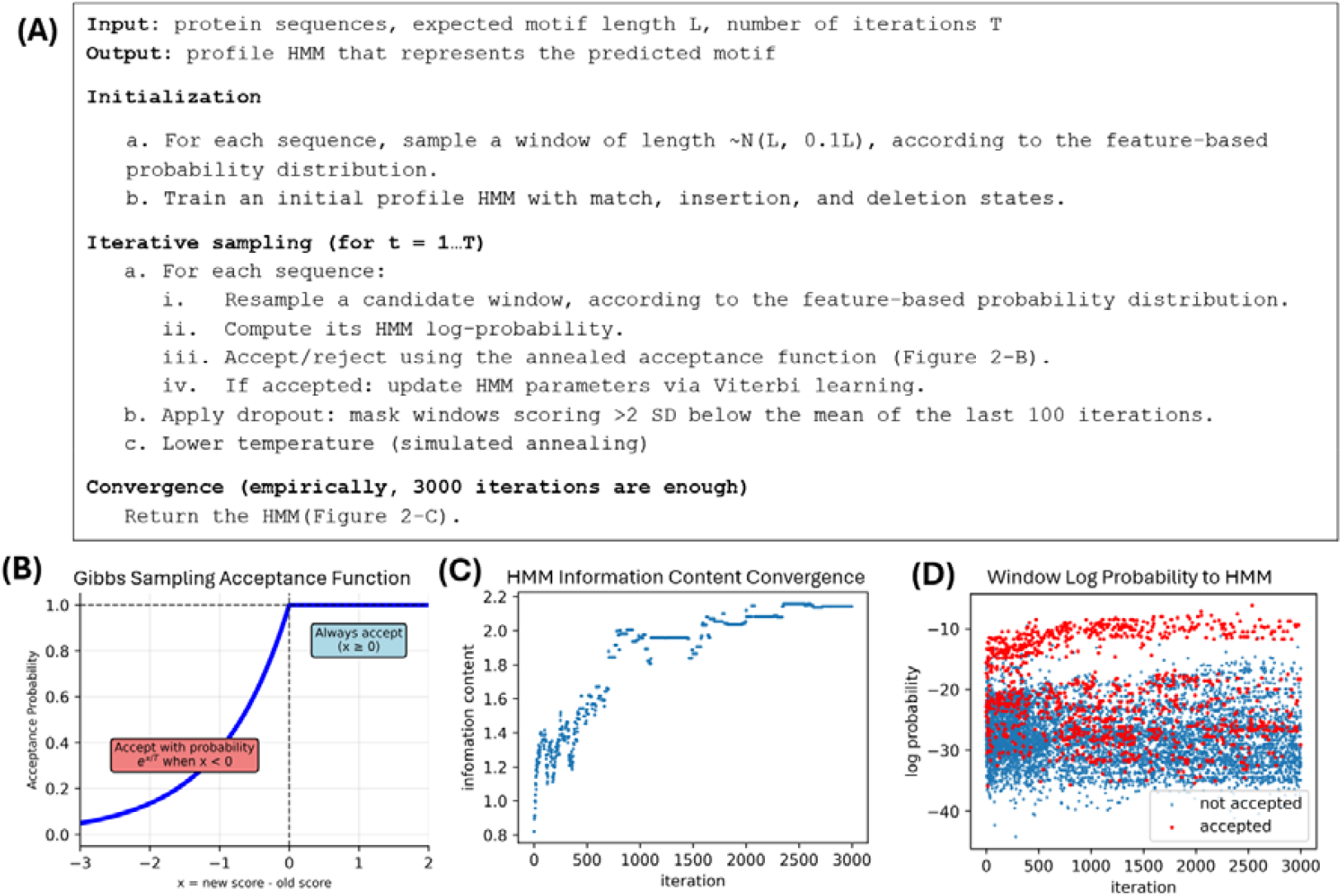
(A) Algorithm for HMM-integrated Gibbs sampling for motif discovery. (B) Acceptance probability function used in our Gibbs sampling algorithm. (C) The increase in information content of the HMM through 3000 iterations. Information content is calculated by averaging the information content of each match state by 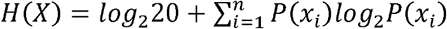. Data generated by running our algorithm on 10 artificial protein sequences. (D) The distribution of log probabilities of all the window samples (10 proteins × 3000 iterations = 30000 samples) using the same setting as (C).

### 2.3 Data preparation

PPI data were obtained from BioGRID v4.4.231 (Oughtred et al., 2021). Tabular files were converted into graph representations using the networkx Python package (v3.1) (Hagberg et al., 2008), with proteins as nodes and PPIs as edges annotated with supporting evidence (throughput, number of publications).

We first exclude all non-human proteins. For each protein, we extracted star networks, defined as the subgraph comprising the protein and all of its partners. Very small star networks (<10 PPIs) were excluded because motifs arising from such limited sets are likely to reflect random similarity rather than meaningful patterns. Very large networks (>200 PPIs) were excluded because they are computationally expensive to analyze and are expected to contain considerable noise. To mitigate this, we applied an additional filter to large networks (>200 PPIs), removing high-throughput interactions not supported by at least two independent publications. Networks falling below the 200-node threshold after this filtering were retained. In total, 12,765 star networks remained for analysis. Networks outside these thresholds can still be constructed and analyzed using the FiGS-MoD scripts available on our GitHub repository. Additionally, a total of 20,419 human protein sequences were obtained from UniProt and used as the reference set (Bateman et al., 2024).

We retrieved 288 motif classes and 2356 human motif instances from the Eukaryotic Linear Motif (ELM) database (Kumar et al., 2024), spanning 1396 human proteins. Each ELM protein sequence was validated against UniProt, and mismatched sequences were excluded. We further noted occasional inconsistencies in the ELM annotation (e.g., motif positions not aligning with the reported residues). To correct these, we re-scanned sequences with the provided ELM regular expressions to ensure positional accuracy. At the end, there were 41 ELM instances not included in our study due to mismatches despite our maximum effort to map them. A reproducible filtering script is available in our GitHub repository for future use as the ELM database expands.

## 3 Results

### 3.1 Simulations

Gibbs sampling theoretically performs best when all input sequences share the same motif class since sequences lacking motifs introduce noise into the model. To evaluate FiGS-MoD under this ideal condition, we input proteins annotated in the ELM database with ARS2 C-terminal leg domain ligand motif (LIG_ARS2_EDGEI_1), SH3 domain ligand motif (LIG_SH3_2), and CtBP ligand motif (LIG_CtBP_PxDLS_1), shown in Figure 3 below. Our method successfully recovered motifs in all three cases. Importantly, in the LIG_ARS2_EDGEI_1 search, we identified a gapped motif containing an indel at position 28 in protein HNRNPLL (Q8WVV9). highlighting the advantage of modeling motifs with an HMM rather than a PSSM. In several cases, predicted motifs did not align exactly with the annotated ELM positions but resembled other motif instances from the same search, suggesting potential novel motif occurrences not yet recorded in ELM.

**Figure 3.**
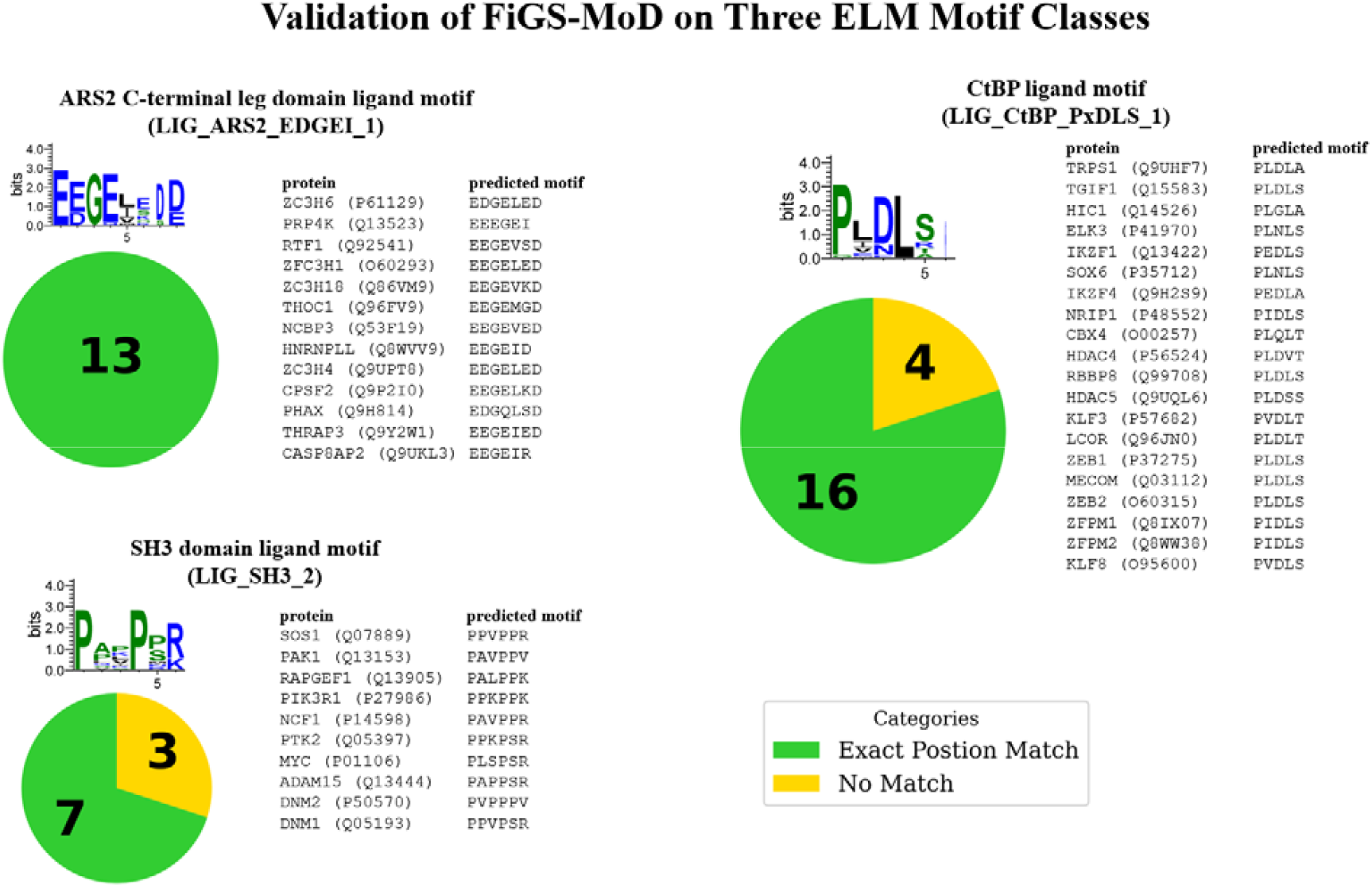
Validation of FiGS-MoD on three known motif classes. Motif discovery was performed for the ARS2 C-terminal ligand motif (LIG_ARS2_EDGEI_1), the SH3 domain ligand motif (LIG_SH3_2), and the CtBP ligand motif (LIG_CtBP_PxDLS_1), using only proteins annotated with these motifs in the ELM database as input. Across all three classes, FiGS-MoD achieved high exact position matches, defined as predictions identical in sequence and location to ELM annotations. Predictions that did not align exactly still showed strong similarity to the ELM class sequence logos, plotted using WebLogo 3.7.9 (Crooks et al., 2004), suggesting the potential identification of additional motif instances not yet catalogued in ELM.

On a separate note, we also tested MEME (v5.5.7) with standard parameters under the zero-or-one occurrence per sequence (ZOOPS) mode on the same input proteins. MEME successfully recovered 13/13 instances of the ARS2 C-terminal ligand motif but did not identify known motifs in the other two datasets. Instead, it reported a consensus motif with the regular expression [FI].C..[CD][GND] for CtBP ligand motif dataset. This illustrates MEME’s strong sequence-driven search capability, though in the absence of biological context, it may also predict enriched sequence patterns that are not necessarily functional motifs.

### 3.2 Case studies

To complement the controlled simulations, we next examined predictions from the full human proteome run. Unlike the simulations, this run analyzed BioGRID star networks (after filtering) to generate the set of predicted motifs in our database. Below, we highlight three case studies using FiGS-MoD.

NCK1 (P16333) is an SH2/SH3 adaptor protein containing three SH3 domains and one SH2 domain (Braverman & Quilliam, 1999). Our algorithm identified several proline-rich motifs in its partners, including KHDRBS1 (Q07666): IPLPPPP, WAS (P42768): IAPPPPT, WIPF2 (Q8TF74): IPPPPPP, WIPF1 (O43516): IPPPVPS, and GAB2 (Q9UQC2): IAPPPRP. These interactions are supported by BioGRID PPIs, and their consensus matches canonical SH3-binding motifs, suggesting they may represent previously unannotated motifs mediating SH3-domain interactions.

In RUNX1 (Q01196), we identified a novel phosphorylation motif predicted to be mediated by MAPK1 (ERK2). The motif exhibited strong structural and evolutionary signatures (disorder >1.0 SD, RSA >1.2 SD, ProtT5 >1.8 SD above background) and the phosphorylation between RUNX1 and MAPK1 was supported by low-throughput experiment previously (Varshney et al., 2012). This case demonstrates both the biological relevance of our prediction and the strength of the discovery framework.

In WEE1 (P30291), we identified a motif in the context of its interaction with SFN protein (P31947). The predicted motif (ARSPTEP), located at residue 64, matches the regular expression R..[ST].P. WEE1 is known to be regulated by phosphorylation (Watanabe et al., 1995), and a phosphosite at this exact position has been reported in a high-throughput study (Ochoa et al., 2020), consistent with our prediction that it mediates WEE1–14-3-3 binding.

### 3.3 Proteome-wide comparison with ground truth

We benchmarked FiGS-MoD against FiGS-MoD without Gibbs sampling and MEME (v5.5.7, standard parameters, ZOOPS mode) on the same star networks. To evaluate performance, we used Fisher’s exact test (FET) to compare each method against a random window sampler baseline by counting overlaps to ELM motifs and PhosphoGRID phosphosites. In this framework, an odds ratio greater than 1 indicates enrichment over random. As shown in Figure 4, FiGS-MoD consistently achieved higher overlap than both FiGS-MoD without Gibbs sampling and MEME, demonstrating improved performance over prior-only and sequence-only discovery methods.

**Figure 4.**
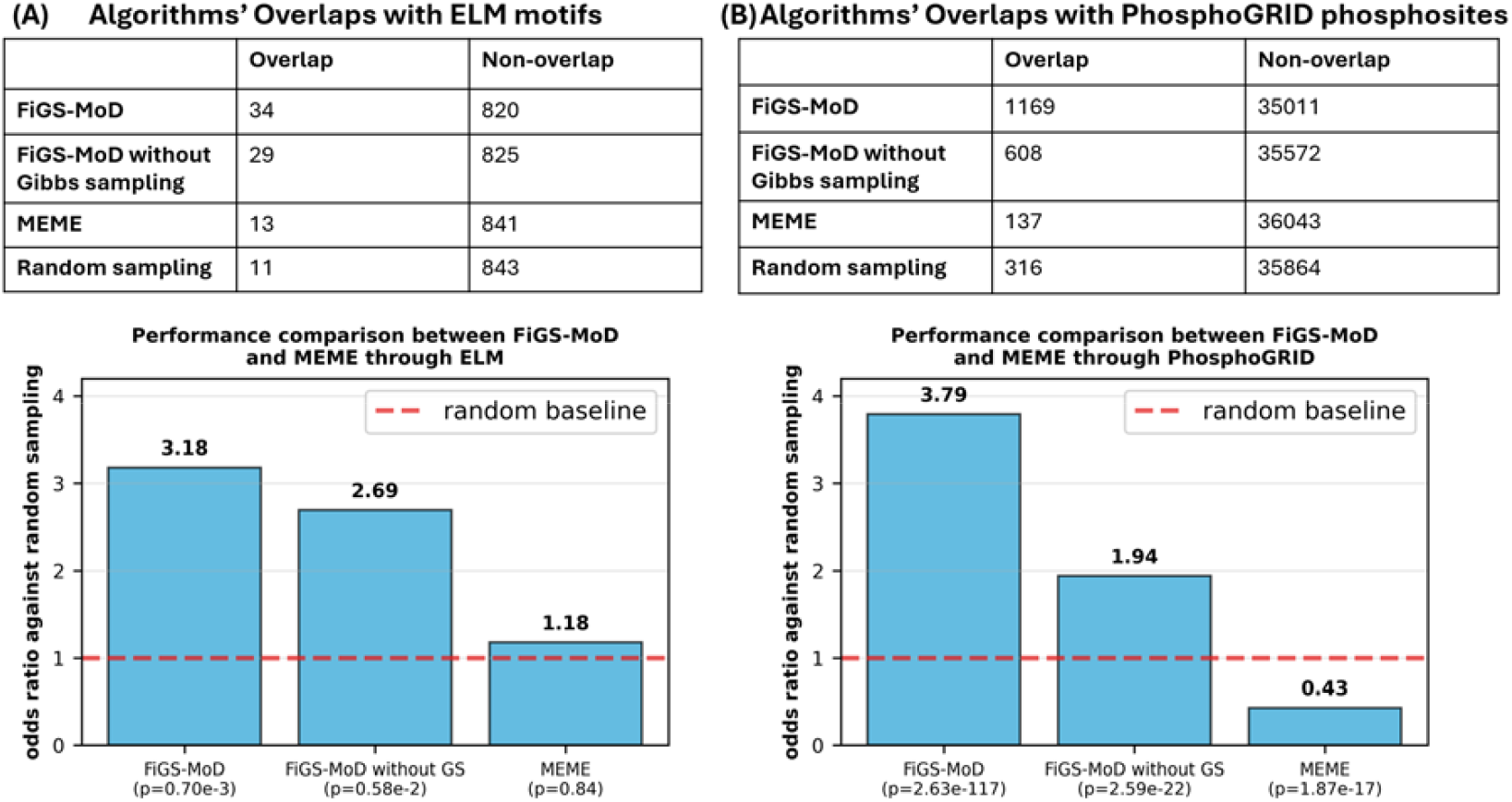
Proteome-wide validation of motif discovery using Fisher’s exact test. Fisher’s exact test (FET) was used to compare the overlap of predicted motifs with annotated ground truths, with each algorithm evaluated relative to a random window sampler baseline on the same star networks. Odds ratios greater than 1 indicate enrichment over random. FiGS-MoD consistently outperformed both FiGS-MoD without Gibbs sampling and MEME in both datasets. (A) ELM validation: restricted to linear motif binding domain (LMBD) star networks where the hub protein contained a binding domain and the partner protein carried the corresponding ELM motif; only proteins for which all three algorithms produced predictions were included. (B) PhosphoGRID validation: restricted to proteins containing annotated phosphorylation sites, again limited to the subset where all three algorithms produced predictions.

For the ELM-based evaluation, we restricted the analysis to predictions consistent with the central assumption of our method: linear motifs interact with hub proteins via binding domains. Specifically, we only considered linear motif binding domain (LMBD) star networks where the hub protein was annotated in ELM as a binding domain-containing protein. We only selected the partner proteins that carry the LMBD corresponding ELM motif as well. To ensure comparability across methods, we further limited evaluation to the subset of proteins for which all three algorithms simultaneously produce predictions, keeping the validation set size the same.

Because ELM motifs were used to train the ProtT5-based residue prior, they do not represent a fully independent validation. To address this, we performed a second evaluation using PhosphoGRID phosphorylation sites, which are independent of our training data but biologically relevant since many phosphosites occur within SLiMs. For this analysis, we only considered proteins with annotated phosphosites and again evaluated the subset where all three algorithms produced predictions. FiGS-MoD significantly outperformed FiGS-MoD without Gibbs Sampling and MEME on this independent dataset, confirming the robustness of our approach.

Finally, comparing FiGS-MoD with and without Gibbs sampling shows that the full algorithm consistently performs better. This indicates that Gibbs sampling provides additional value beyond residue-level priors, refining candidate motifs and improving overlap with validated sites.

### 3.4 Contribution of PLM

To examine which feature scores drive residue annotation, we compared the distributions of four features: ProtT5 language model (PLM scores), disorder, RSA, and conservation, across motif residues, domain residues, and other residues (Figure 5). An effective feature should clearly separate motif residues from the other categories. Among the four, PLM exhibited the strongest discrimination. Disorder and RSA provided weaker but complementary signals; for example, disorder distinguished motifs from domains, discouraging false positives within structured regions. Conservation separated motifs from other residues only modestly, but the difference remained statistically significant (p = 1.37 × 10□□□, motif versus neither).

**Figure 5.**
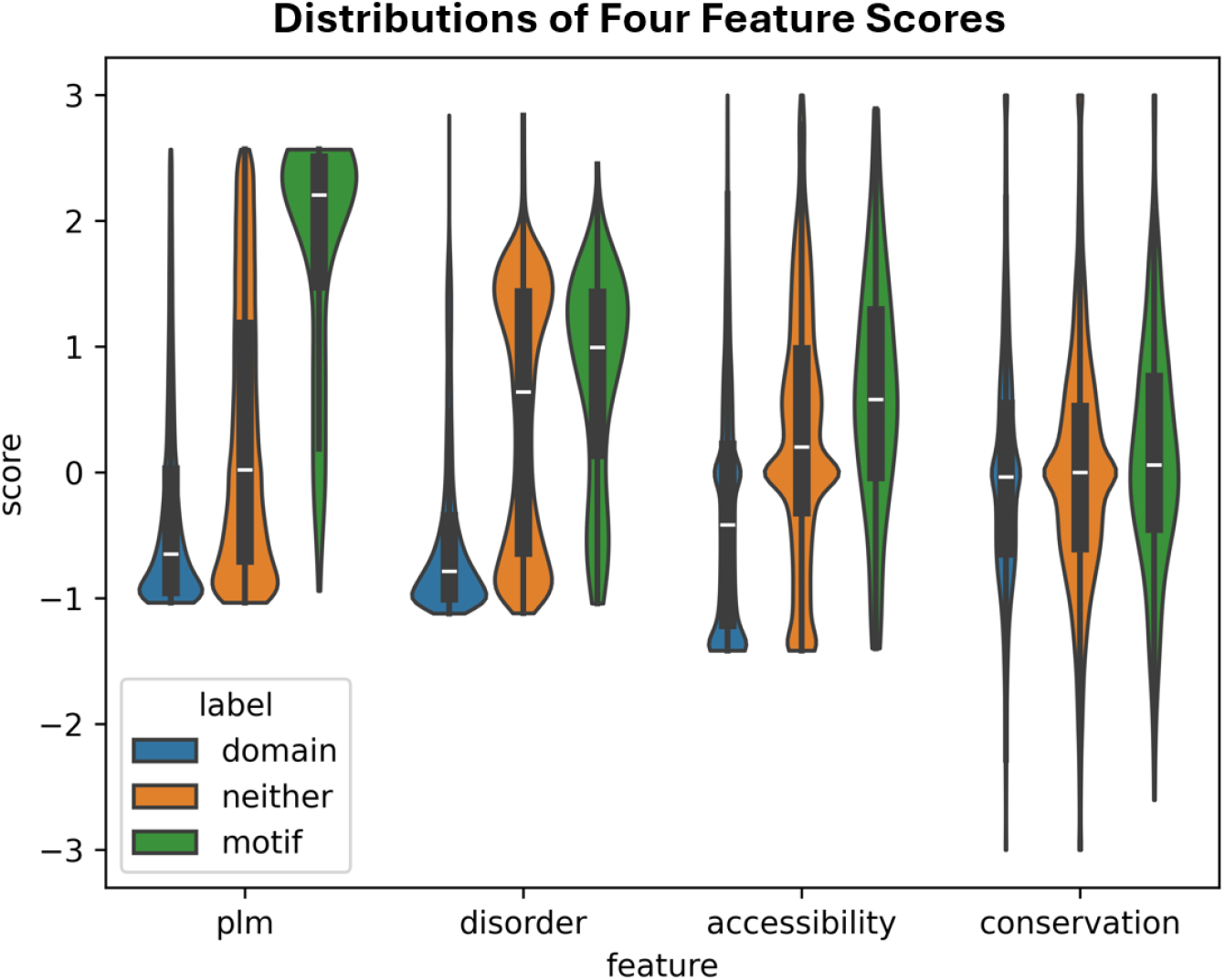
Contribution of residue-level features to motif annotation. Violin plots show the distributions of four z-normalized feature scores for 500,000 sampled residues, divided into motif, domain, and non-motif/non-domain (neither) categories. The protein language model (PLM; ProtT5) score provides the strongest separation between motif and non-motif residues, while disorder, relative solvent accessibility (RSA), and evolutionary conservation contribute complementary signals (e.g., disorder helps distinguish motifs from domains; conservation integrates ortholog information). These distributions highlight which features most effectively guide FiGS-MoD in identifying biologically meaningful motifs.

## 4 Discussion

The ProtT5 language model is central to our framework. The residue-level scores we derived from its embeddings effectively distinguish motif from non-motif residues (Figure 5), providing a strong baseline signal. Additional features, such as disorder, RSA, and conservation, complement ProtT5 by capturing information the language model alone cannot, including evolutionary context. While PLM (supervised learning) profiling effectively prioritizes candidate residues, Gibbs sampling (unsupervised learning) leverages information content from interactomic data to refine motif discovery within through a heuristic framework. Together, these features yield improved predictions, and the combination of PLM priors with Gibbs sampling outperformed either component alone. More broadly, this illustrates the value of integrating supervised learning with unsupervised discovery in biological problems where curated labels are sparse relative to the true number of functional instances, as in linear motif discovery.

Two main limitations remain. First, Gibbs sampling is inherently stochastic and may yield different outputs under different random seeds. Despite our efforts to reduce this variability through residue-level priors and dropout filtering, the nondeterministic nature of the method cannot be fully eliminated. Importantly, this also represents an opportunity: running FiGS-MoD multiple times or under alternative parameter settings can reveal additional motif instances that may not appear in a single run. In this sense, the algorithm’s predictions are not yet exhausted, and users are encouraged to experiment with repeated runs or extended iterations on their own protein datasets, which may uncover additional motif instances beyond those identified currently.

Second, computational validation of predicted motifs remains challenging. Only ∼0.2% of the estimated SLiMs, which are about one million in total, in the human proteome are currently annotated in the ELM database (Tompa et al., 2014), leaving uncertainty as to whether predictions outside of ELM are false positives or represent novel, uncharacterized motifs. Ultimately, experimental validation is required. However, high-throughput techniques for validating transient motif-mediated PPIs remain limited. Emerging methods such as cross-linking mass spectrometry may provide a promising avenue for systematic validation in the future (O’Reilly & Rappsilber, 2018; Veale & Clarke, 2024).

